# Guided by touch: Tactile Cues in Hand Movement Control

**DOI:** 10.1101/2024.07.26.605248

**Authors:** Maria Evangelia Vlachou, Juliette Legros, Cécile Sellin, Dany Paleressompoulle, Francesco Massi, Martin Simoneau, Laurence Mouchnino, Jean Blouin

## Abstract

Traditionally, touch is associated with exteroception and is rarely considered a relevant sensory cue for controlling movements in space, unlike vision. We developed a technique to isolate and evaluate tactile involvement in controlling sliding finger movements over a surface. Young adults traced a 2D shape with their index finger under direct or mirror-reversed visual feedback to create a conflict between visual and somatosensory inputs. In this context, increased reliance on somatosensory input compromises movement accuracy. Based on the hypothesis that tactile cues contribute to guiding hand movements, we predicted poorer performance when the participants traced with their bare finger compared to when their tactile sensation was dampened using a smooth finger splint. The results supported this prediction. EEG source analyses revealed smaller current in the presumed somatosensory cortex during sensory conflict, but only when the finger directly touched the surface. This finding suggests the gating of task-irrelevant somatosensory inputs. Together, our results emphasize touch’s involvement in movement control, challenging the notion that vision predominantly governs goal-directed hand or finger movements.

## Introduction

The sense of touch, deeply rooted in our evolutionary history, has evolved into a sophisticated and versatile sensory modality. Once imperative for navigating and guiding animal behavior^1^, touch has, in the modern human experience, moved beyond its immediate necessity for survival. The ability to engage in human activities like Braille reading and touch screen interactions implies that touch surpasses a mere interface with the external world, hinting at the preservation of its primitive function in guiding spatially oriented movements during primate phylogenesis. Supporting this assumption are the striking similarities in the tactile-based somatosensory topographical organization of the midbrain, a crucial region for orienting movements, in both reptiles (tectum) and mammals (superior colliculus)^2^.

Tactile feedback, transmitted through our fingers’ skin, is a language of its own. Not only does it provide a plethora of details about the objects we interact with, such as their texture and shape^3,4^, but it also decodes information about the body position and motion, contributing to proprioception^5–7^. The pivotal observation by Hulliger et al.^8^ that skin mechanoreceptor activity undergoes changes during isotonic finger movements performed without direct physical contact was fundamental in conceptualizing touch as a conveyor of movement-related cues. Behavioral experiments further substantiated the link between touch and movement perception by demonstrating that the mere stretching of the skin elicits illusions of movement in the joints covered by the skin^6,9,10^.

Transitioning to a neurophysiological perspective, nerve recordings in healthy subjects captured the sensitivity of mechanoreceptors in encoding movement direction^11^. Similarly, investigations in monkeys have unveiled differential responses of Merkel discs (SAI) and Meissner corpuscles (FAI) afferents to tangential forces applied in different directions^12^. Importantly, neurons with direction selectivity abound within the somatosensory cortex^4,13^. Together, these receptors and neurons may constitute a critical physiological substrate underpinning our capacity to discern the displacement of our hand in relation with external surfaces^14,15^.

Recent studies showed that, without hand visual feedback, the direction of voluntary sliding finger movements on a surface with parallel ridges is altered by the ridge’s orientation^16,17^. This observation extends our understanding of touch beyond exteroception, highlighting its contribution to controlling the orientation of finger movements on touched surfaces. However, whether this tactile motor function still holds true when visual feedback on hand location and movement is available remains unknown. It is imperative to recognize that in most real-world scenarios, visual feedback profoundly shapes our sensorimotor interactions, emphasizing the need to consider multisensory integration processes when evaluating the motor function of touch. Indeed, the remarkable flexibility of the brain, its ability to dynamically control the transmission of afferent signals^18–20^, and the reliability of visual cues in controlling movements^21^ converge to suggest that tactile involvement in motion control might be limited solely to contexts where hand vision is precluded, as was the case in Moscatelli et al.^16^ and Bettelani et al.^17^. The increased responsiveness of the somatosensory cortex to foot tactile stimulation observed in individuals with impaired vestibular systems^22^ supports the hypothesis of an upregulation of relevant tactile inputs in sensory impoverished contexts.

The challenge of investigating whether skin sensory input contributes to the control of hand movement in the presence of visual feedback arises from the rapid ceiling effect observed in visually-guided tasks, which obscures the genuine impact of tactile input. At the neurophysiological level, inferring cutaneous contributions from the activity of the primary somatosensory cortex (S1) in humans is challenging. This challenge primarily stems from the proximity of distinct tactile and proprioceptive representations in the somatotopic map^23^, as well as the presence of cell populations that encode both sensory modalities^24–26^. We employed an established task to assess the extent to which somatosensory inputs are involved in the sensory-motor loop controlling movements. Specifically, healthy young human adults traced with their index finger the outline of a two-dimensional shape while receiving mirror-reversed visual feedback about their movements (Methods, Fig. 1). This context elicits a conflict between afferent visual and somatosensory cues^27^, and movements become inaccurate and jerky^20,28,29^. However, studies have shown that reducing somatosensory inflow to the cerebral cortex, whether through neuropathies^29^, inhibition of the somatosensory cortex via repetitive transcranial magnetic stimulation^30^, or dynamic somatosensory gating^20^, results in reduced movement perturbation under mirror-reversed vision.

**Fig. 1.**
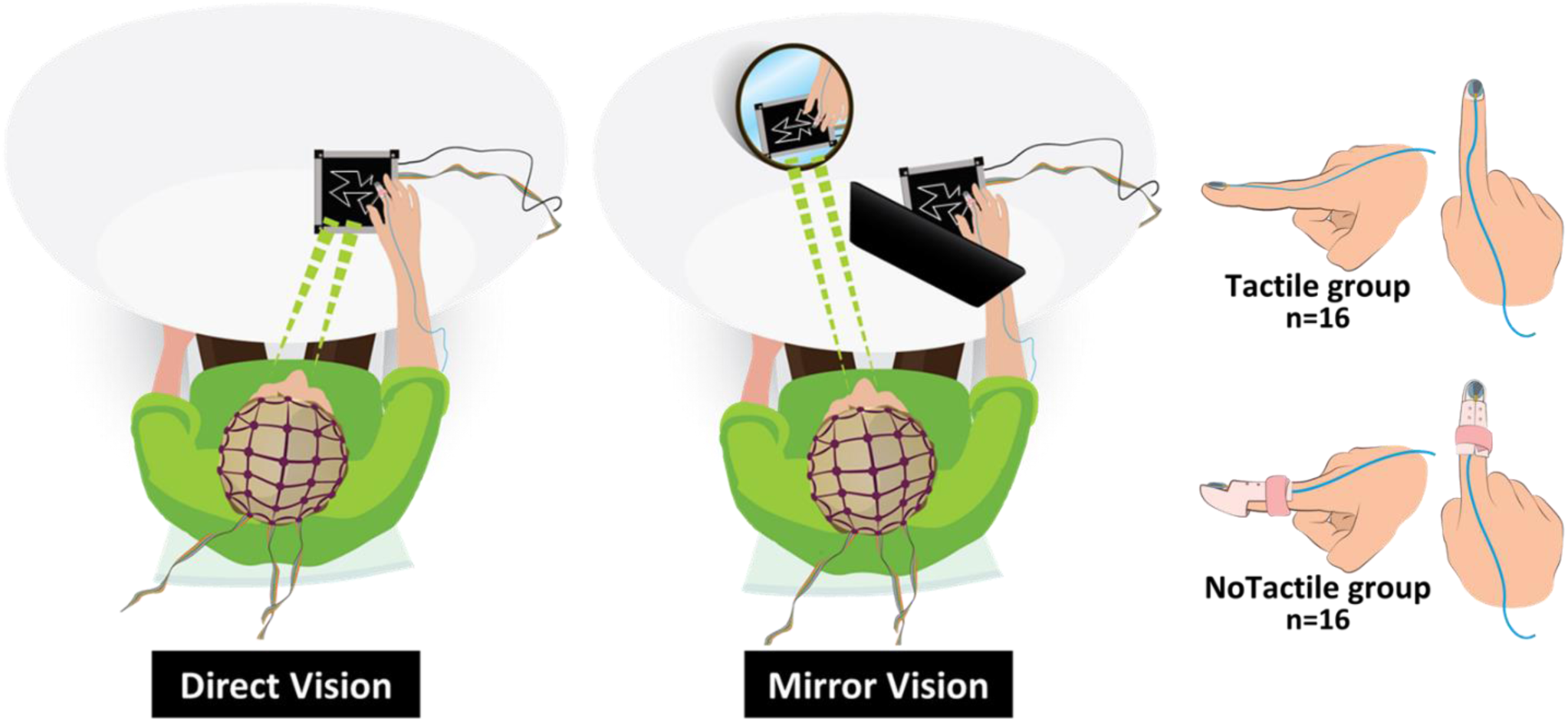
Experimental protocol. Participants slowly traced the outline of a polygon printed on a textured surface with their index finger while looking both at the polygon and their hand either directly (Direct condition) or through an inclined mirror (Mirror condition). In the Mirror condition, a black shield blocked the direct view of their hand during tracing. Participants of the Tactile group directly touched the surface with the pulp of their index finger. Participants of the NoTactile group wore a plastic finger splint to reduce tactile stimulation from the surface. In both groups, an accelerometer fixed on the right index finger’s nail estimated the vibratory tactile stimuli during tracing, and a force platform recorded the X-Y-Z forces exerted on the surface.

Here, we tested the hypothesis that tactile feedback contributes to guiding spatial movements by instructing participants to trace the contour of a shape with their index finger in a conflicting visuo-somatosensory environment. Our rationale posits that if tactile inputs participate in guiding hand movements, tracing performance will be less affected by mirror-reversed visual feedback when tactile feedback is reduced. To achieve this tactile reduction, participants wore a smooth, rigid finger splint (Fig. 1). Furthermore, we anticipated that during the intersensory conflict, the somatosensory cortex would show reduced activity (i.e., sensory gating) when tactile inputs provide information on the relative finger-surface motion (i.e., bare index finger) as compared to when these inputs are diminished using a finger splint. This hypothesis builds upon findings suggesting that the central nervous system can exert dynamic control over the transmission of task-irrelevant afferent signals^20,31,32^.

## Results

### Tactile stimulation

Participants slowly (velocity ∼10 mm/s) traced the outline of a polygon printed on a textured surface (See Methods Fig. 6) with their index finger while looking both at the polygon and their hand either directly (Direct condition) or through an inclined mirror (Mirror condition). Participants of the NoTactile group wore a plastic finger splint to reduce tactile stimulation from the surface. In contrast, participants of the Tactile group directly touched the surface with the pulp of their index finger (Fig. 1). The pulp of the index finger is densely packed with mechanoreceptors, containing approximately 40% of the total mechanoreceptors found in the entire finger^33^. This high concentration of mechanoreceptors makes the fingertip exceptionally sensitive and adept at detecting fine textures, vibrations, and minute changes in surface properties, thereby making it particularly well-suited for guiding the hand across surfaces. The transient mechanical stimuli generated by the sliding finger movement on the surface mainly targeted the skin receptors that are activated by vibration (e.g., SAI and FAI) and skin stretches (e.g., SAII). To test if the use of finger splint genuinely attenuated tactile stimulation, we compared, between the Tactile group (bare finger) and the NoTactile group (with finger splint): a) the Power Spectral Density (PSD) of the acceleration as measured from an accelerometer glued on the index finger’s nail, b) the square root of the integrated PSD signal beyond 10 Hz to quantify the amplitude of the acceleration (i.e., vibrational stimuli) and c) the Coefficient of Friction (COF) calculated from the normal (Fz) and tangential (Fx and Fy) contact forces exerted on the surface (see Methods).

Figure 2a depicts the mean PSD of acceleration of all participants of the Tactile and NoTactile groups, under the Direct and Mirror vision conditions. The resulting signal is composed of a large frequency spectrum for both experimental groups. We calculated the integral of the PSD signal of the vertical acceleration within the 5 to 45 Hz frequency range, specifically targeting the response frequencies of tactile SAI and FAI afferents. These mechanoreceptors are involved in tactile sensation, particularly in detecting low-frequency pressure, indentation, and vibration^34^. Critically, SAI and FAI receptors also exhibit directional sensitivity^12^, making them well-suited for controlling finger movement orientation during the tracing task. A 2 (Group: Tactile, NoTactile) × 2 (Vision: Direct, Mirror) mixed ANOVA (alpha level set at 0.05), with Vision as a repeated factor, revealed a significant group difference. The NoTactile group displayed lower integrated PSD values than the Tactile group (F_1,27_=11.35, p=0.002, partial η^2^=0.3), indicative of reduced power within the 5-45 Hz frequency range.

**Fig. 2.**
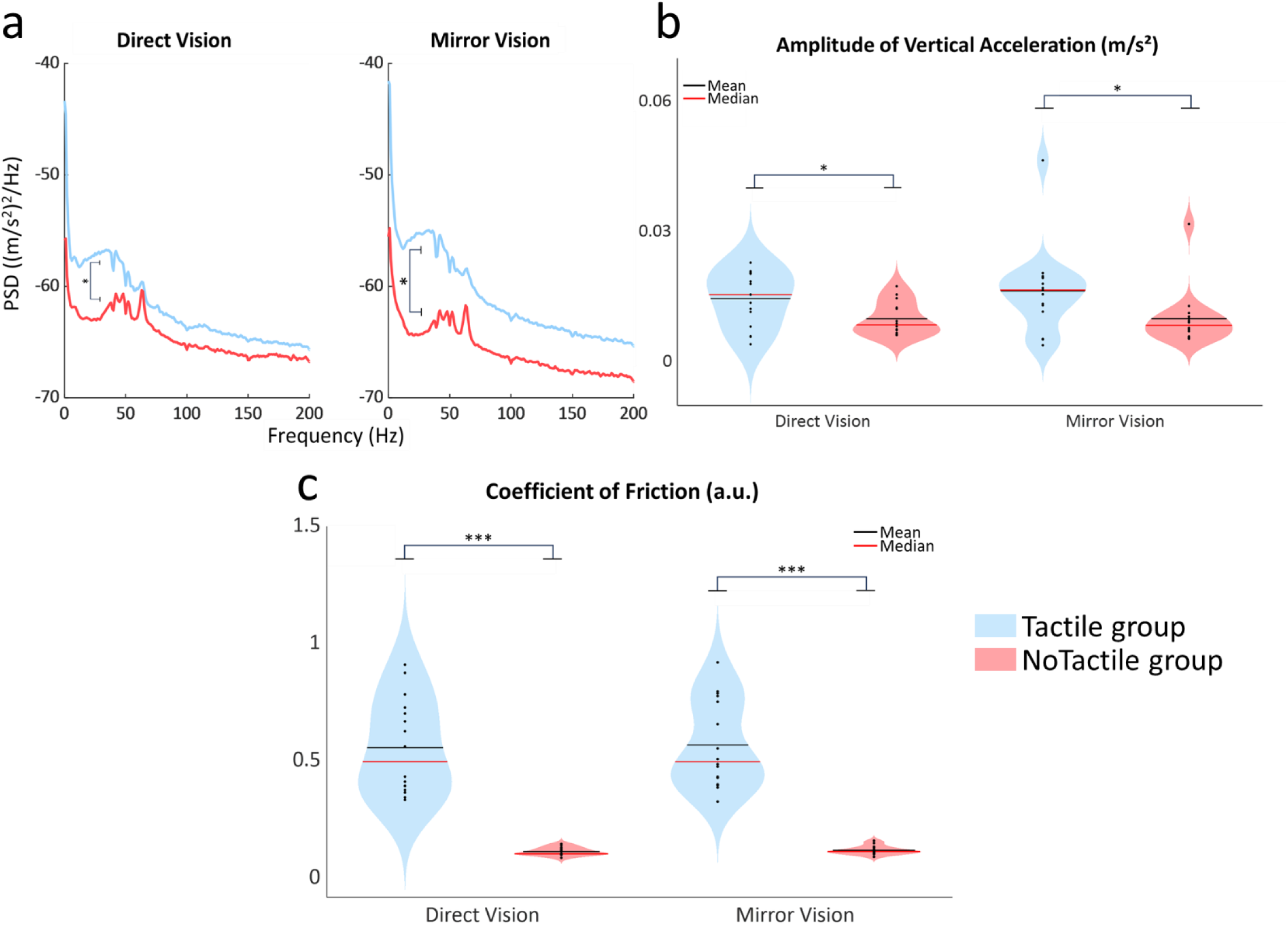
Tactile stimulation is attenuated when direct contact with a surface is interrupted with the use of a finger splint. **a** Mean PSD signal of the vertical acceleration across all trials. The 5-45 Hz frequency range was used for the ANOVA analysis. **b** Violin plots of the square root of the integrated PSD signal beyond 10 Hz, serving as an index of the amplitude of vertical acceleration, during Direct and Mirror tracing. **c** Violin plots of the mean coefficient of friction of all the trials during Direct and Mirror tracing. In panels b and c, individual data points shown in each violin plot represent the mean computed for each participant. For all panels, Blue: Tactile group. Red: NoTactile group. *: p < 0.05, ***: p < 0.001.

A 2 (Group: Tactile, NoTactile) × 2 (Vision: Direct, Mirror) mixed ANOVA revealed that the square root of the integrated PSD signal was significantly lower in the NoTactile group than in the Tactile group (F_1,27_=6.38, p=0.017 partial η^2^= 0.19) on (Fig. 2b). Within each group, one participant exceeded the threshold conventionally accepted to be considered as outliers for data following a normal distribution (i.e. current ±3 standard deviations from the mean^35^). These participants were excluded from the statistical analysis.

The NoTactile group also exhibited lower COF in both visual conditions compared to the Tactile group (Fig. 2c) as evidenced by the significant effect of the experimental group revealed by a 2 (Group: Tactile, NoTactile) × 2 (Vision: Direct, Mirror) mixed ANOVA (F_1,29_=83.93, p<0.00001, partial η^2^= 0.74). There was no significant effect of Vision or Vision x Group interaction, indicating that friction levels decreased with the use of the finger splint and were unaffected by the visual conditions.

Overall, the analyses of the vibration measured on the index finger and of the coefficient of friction, evoked by the finger surface interaction, are consistent with greater tactile stimulation when the participants traced the shape with their bare index finger compared to when they wore a finger splint.

### Behavior: Tracing performance during mirror-reversed visual feedback is more perturbed with unaltered skin-surface tactile feedback

Tracing performance was evaluated by considering the jerk index, computed here by deriving, as a function of time, the tangential contact forces on the textured surface. Jerk measures the smoothness (or “abruptness”) of changes in applied force and is a key variable for assessing sensorimotor impairments^36^. In protocols using conflicting sensory environments, the derivative of contact forces can effectively highlight movement intermittencies that are not easily captured by kinematic data, such as during freezing-like episodes that frequently occur when tracing with mirror-reversed visual feedback and that do not result in tracing errors. Higher values of the jerk index indicate more abrupt and erratic movements, while lower jerk index values reflect smoother and more controlled movements.

The average sum of the calculated absolute jerk index, computed for each participant in the Direct and the Mirror conditions, is shown in Fig. 3 for both the Tactile and NoTactile groups (see Supplementary Fig. S1 and Table S4 for the evolution of jerk index across trials). A 2 (Group: Tactile, NoTactile) × 2 (Vision: Direct, Mirror) mixed ANOVA analysis revealed a significant Group × Vision interaction (F_1,29_=6.45, p = 0.017, partial η^2^=0.18). Post-hoc analyses (Newman-Keuls, p < 0.05) indicated that the jerk index was significantly higher in the mirror condition only within the Tactile group. The lack of a significant difference in the jerk index between the Mirror and Direct conditions for the NoTactile group (p = 0.82) suggests that tracing performance remains largely unaffected by mirror-reversed visual feedback when finger tactile inputs are reduced. Subsequently, these findings serve as a behavioral basis for comparing the cortical activity between the two experimental groups when confronted with a visuo-somatosensory conflict.

**Fig. 3.**
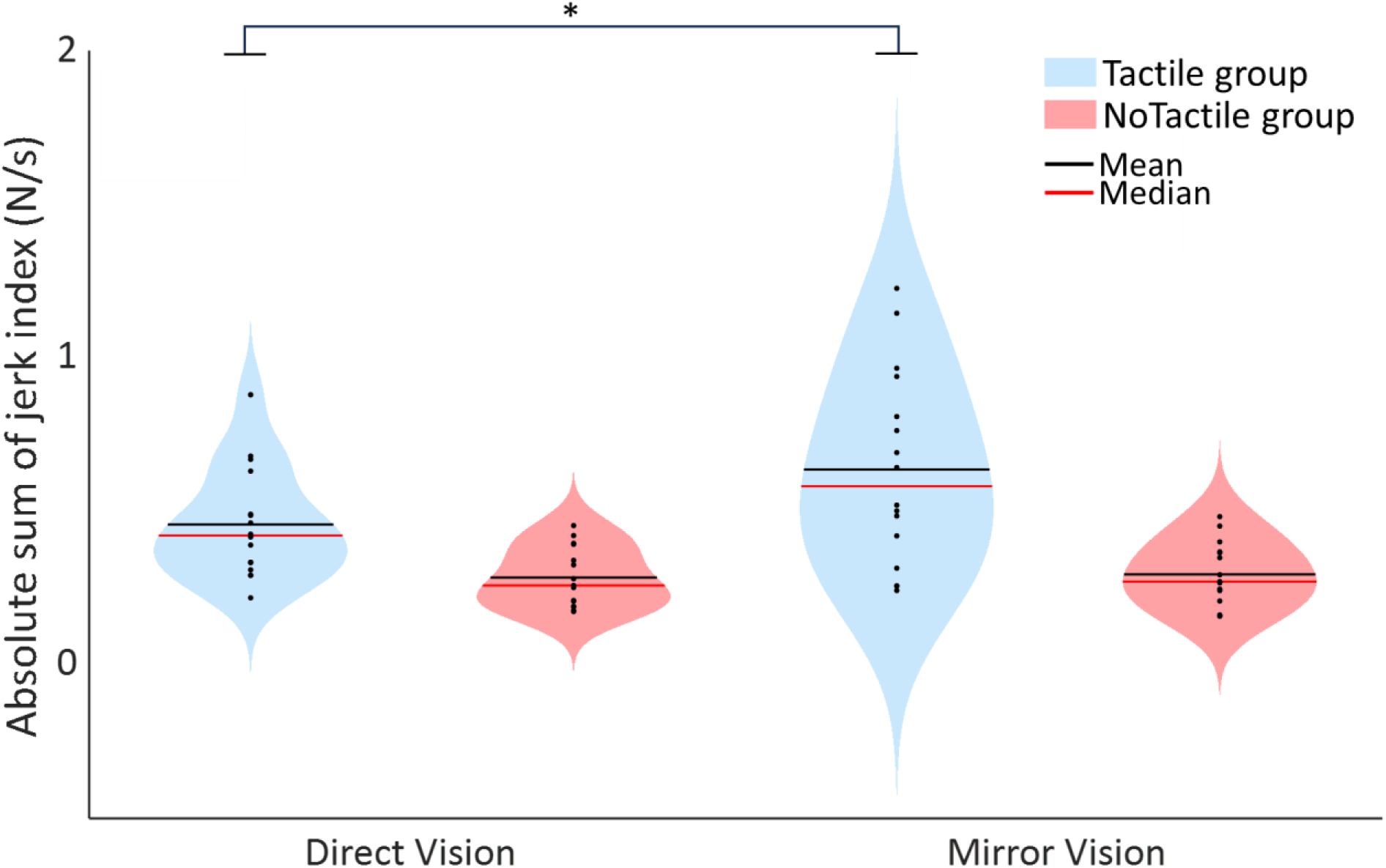
Movements become jerkier during mirror tracing when there is contact between the finger’s skin and the surface. Violin plots of the mean sum of absolute jerk index of all trials for the Tactile (blue) and the NoTactile group (red) during Direct and Mirror tracing. Individual data points represent the mean sum of jerk index computed for each participant and are represented with scatter plots within each violin plot. *: p < 0.05.

### EEG data: Gating of somatosensory inputs if there is contact between the finger’s skin and the surface when tracing with mirror-reversed visual feedback

In the source space, we estimated the mean absolute amplitude of EEG current during the tracing phase to gauge cortical activity^37,38^. Employing parametric paired t-tests (significance threshold p < 0.05, uncorrected), we contrasted the current between the Mirror and the Direct conditions for both the Tactile and NoTactile groups. In the resulting statistical-topographical current maps shown in Fig. 4, warm colors indicate greater cortical activation in the Mirror relative to the Direct condition, while cool colors represent diminished cortical activation. Therefore, when observed in sensory areas, cool colors suggest sensory gating of the sensory feedback, while warm colors reflect somatosensory facilitation.

**Fig. 4.**
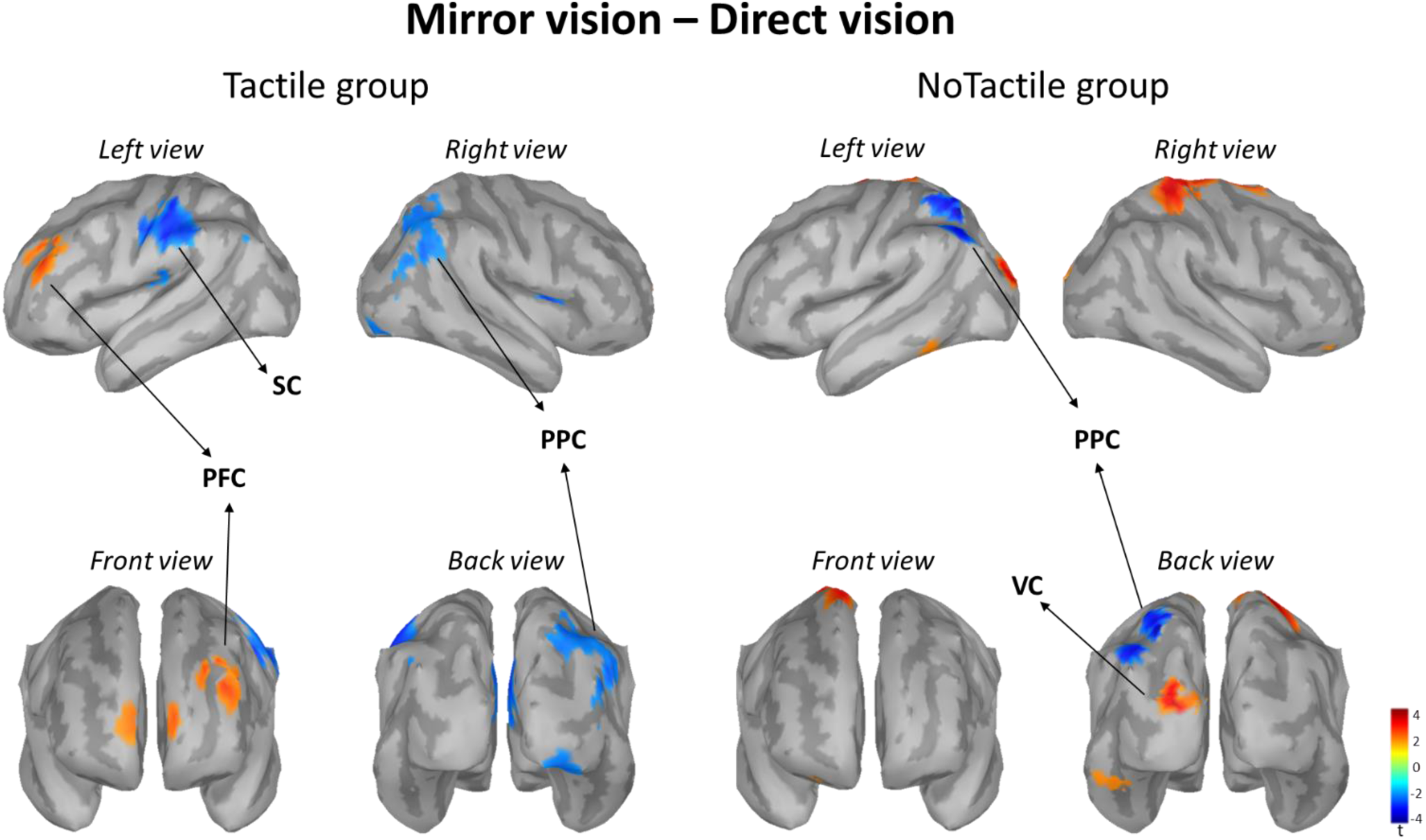
Statistical maps of the absolute current amplitude in the source space resulting from the contrast between Mirror condition and Direct condition for both Tactile (left) and NoTactile (right) groups. Sources are displayed on a cortical template (MNI’s Colin 27). Side (left and right), front and back cortical views are displayed for each contrast.

For both the Tactile and NoTactile groups, significant differences in the estimated cortical current emerged between the Mirror and the Direct conditions. In the Tactile group, these differences manifested notably in the presumed contralateral somatosensory cortex (SC) to the drawing hand (i.e., left hemisphere) with decreased neuronal activity while tracing with incongruent visuo-somatosensory inputs. This observation suggests a gating of somatosensory information with mirror-reversed vision in presence of movement-related tactile inputs. A reduced activity was also observed in the right posterior parietal cortex (PPC), whereas an increased activity was evident in the left prefrontal area (PFC). These fronto-parietal activity changes may have played a key role in reducing the weight of tactile cues in sensory integration processes (see discussion).

The NoTactile group exhibited no significant difference in the current amplitude over the left SC between the two visual conditions. However, the mirror vision led to a significant increased current in the left visual cortex (VC) and a significant decreased current in the left PPC.

Our main prediction was that the Tactile group would exhibit reduced activity in the somatosensory cortex while tracing the shape with mirror-reversed vision compared to the NoTactile group. To test this hypothesis, we computed for each participant the mean absolute current amplitude recorded during the tracing over the left postcentral gyrus region of interest (ROI), as topographically defined in Destrieux^39^ and shown in Fig. 5. As supported by a significant Group (Tactile, NoTactile) × Vision (Direct, Mirror) interaction (F_1,27_=4.6, p=0.04, partial η^2^= 0.18), the activity of the somatosensory cortex decreased for the participants of the Tactile group when tracing with mirror-reversed feedback (p=0.01) but did not significantly differ between the two visual conditions for the NoTactile group (p=0.8) (Fig. 5). Within each group, the EEG data of one participant exceeded the conventionally accepted threshold for outliers and were excluded from the ROI analyses (see Supplementary Fig. S2 & Fig. S3).

**Fig. 5.**
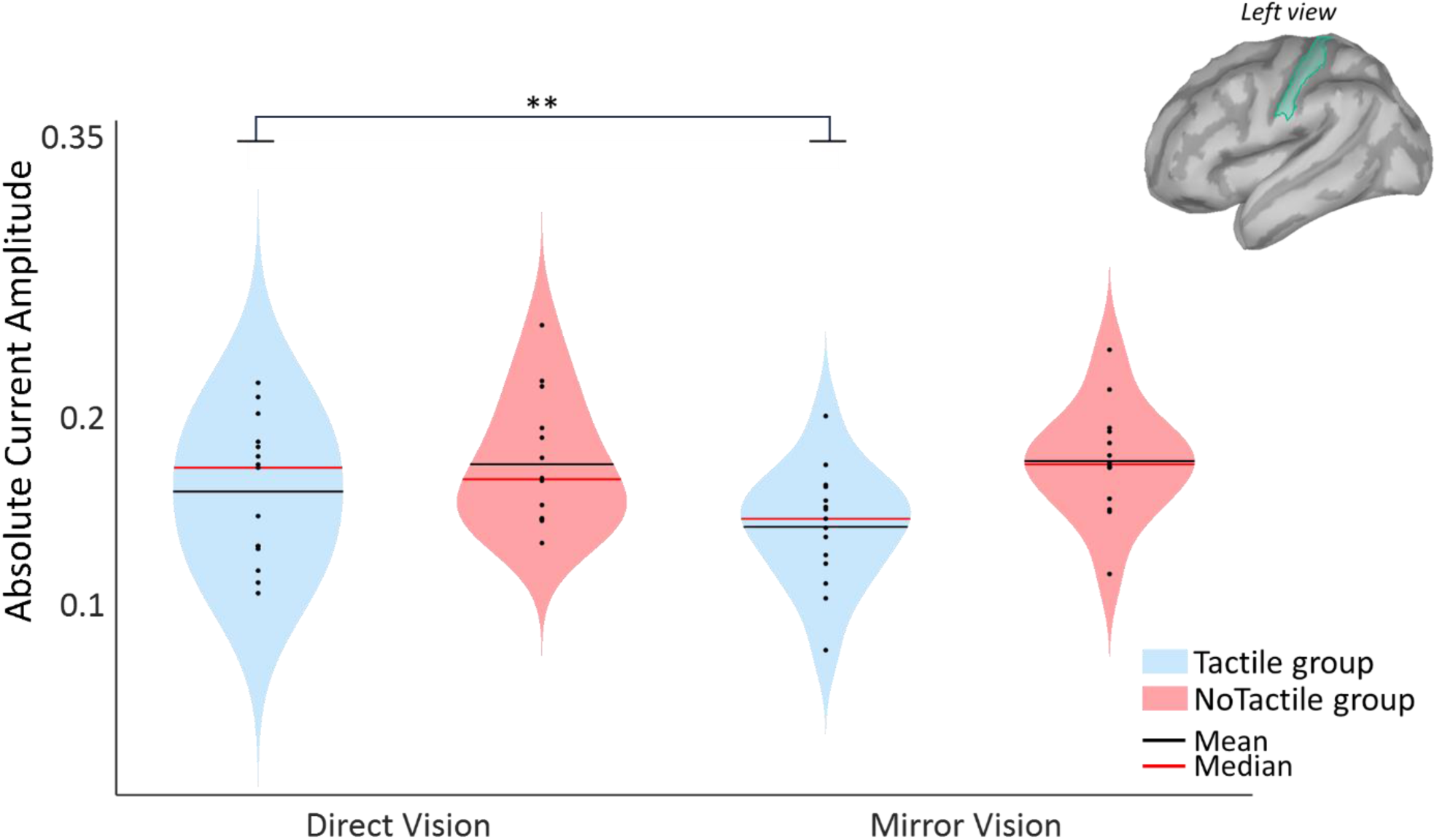
Decreased activity in the somatosensory cortex with mirror-reversed vision if there is contact between the finger’s skin and the surface. Violin plots of the mean absolute current computed over the left postcentral gyrus ROI, as topographically defined in Destrieux^39^ (depicted in the right inset), during the Direct and Mirror conditions for both Tactile (blue) and NoTactile (red) groups. Individual data points represent the mean current computed for each participant and are depicted with scatter plots within each violin plot. **: p < 0.01.

## Discussion

The present study delves into the intricate interplay between tactile feedback and voluntary movements, specifically focusing on motor tasks incorporating spatial orientation. While previous studies have substantiated the coupling of touch and motion perception^14,15^, the question of whether the brain uses skin-derived afferent information to control spatially oriented movements remains largely unexplored. Recent work by Moscatelli and colleagues^16^ tackled this question, providing evidence of tactile involvement in the spatial guidance of finger movements against a surface in the absence of visual feedback. Importantly, the blindfolded approach used in their study may have altered the sensory weighting process of visuo-somatosensory signals, potentially attributing greater relative weight to the sense of touch compared to conditions where visual feedback is available. This possibility is supported by studies showing that sensory deprivation can potentiate spared sensory inputs, a phenomenon known as sensory substitution^22,40–43^.

Participants engaged in a tracing task, following the contour of a shape with their index finger while receiving either direct or mirror-reversed visual feedback of their movements. Mirror vision creates a visuo-somatosensory conflict, and previous studies have shown that in this novel sensory context, increased processing of somatosensory input leads to poorer tracing performance^20,29,30^. By comparing the motor behavior of participants wearing a finger splint that reduces tactile sensation (NoTactile group) with that of participants receiving normal tactile feedback during finger sliding movements (Tactile group), we were able to elucidate whether touch plays a role in hand movement control. The efficacy of the splint as a tactile attenuator during surface scanning was confirmed by the amplitude and power spectral density (PSD) of finger vertical acceleration that were both significantly reduced compared to the Tactile group, as well as the reduced COF exhibited by the NoTactile group. These results indicate that less energy was transferred through the cutaneous receptors in the NoTactile group, leading to reduced sensory processing regarding the skin’s stress state. Considering these key physical variables related to skin stimulation, we confidently assert that the NoTactile group, using the finger splint, experienced diminished tactile stimulation while tracing the shape’s contour compared to the Tactile group, who directly touched the surface with their index finger.

Remarkably, mirror-reversed vision feedback disrupted tracing performance only for participants in the Tactile group. Specifically, when tracing the shape with their bare finger, these participants exhibited significantly jerkier movements under mirror vision compared to direct vision. This reduction in movement smoothness highlights the participants’ heightened susceptibility to the sensory conflict introduced by the mirror, impairing their control over tracing movements. In contrast, the movement smoothness profile of participants in the NoTactile group did not differ significantly between the mirror and direct vision conditions. This consistent performance suggests reduced sensory conflict with diminished tactile feedback.

Performing the tracing task with the bare index finger, as opposed to using a finger splint, likely introduced an additional sensory channel (tactile) that, along with the proprioceptive channel, provided information related to the displacement of the finger on the surface. The prevailing explanations for motor impairments observed when visual feedback is distorted have primarily focused on the sense of proprioception^20,44–46^, overlooking the potential contribution of cutaneous afferents. In the present study, we demonstrated that enhancing the somatosensory input with tactile information from the finger-surface interaction increased visuo-somatosensory conflict, leading to more abrupt and corrective actions of the tracing hand. Therefore, the significant contribution of tactile feedback in spatially controlling sliding finger movements, as evidenced by Moscatelli et al.^16^ in the absence of visual feedback, persists when vision is available.

It has been suggested that the dynamic suppression of cortical somatosensory inputs observed in individuals performing goal-directed movements under conflicting visuo-somatosensory feedback serves to partially resolve the sensory conflict^20,47,48^. This sensory gating was observed in the Tactile group, but not in the NoTactile group. This suggests that the sensory gating phenomenon primarily targeted tactile rather than proprioceptive feedback. Proprioception plays a crucial role in motor control by providing the brain with information about the position, orientation, and movement of various body parts, including the fingers^49^. Previous studies exploiting the mirror paradigm reported gating over the somatosensory cortex even when participants traced a shape with the tip of a pen^20,48^. In this light, one might question why the tracing performance and the activity of the source-localised somatosensory cortex were unaffected by the mirror-reversed visual in the NoTactile group. Several pieces of evidence suggest that the gating of the somatosensory cortex observed in previous studies may have been intricately linked to tactile cues. When tracing with a pen, hand tactile receptors are stimulated by the pressure of skin with the pen, the skin-pen relative motion, and by the vibration transmitted by the sliding pen and the surface. Research has shown that eliminating this information can affect one’s ability to precisely control the movement of a pen on a surface^50^. Therefore, tactile input provides information about pen motion and becomes irrelevant in contexts with visuo-somatosensory conflict. This observation could explain the somatosensory gating reported in previous studies, where participants traced the contours of shapes with the tip of a pen.

Another plausible explanation is that proprioceptive feedback was diminished in the NoTactile group by the use of a finger splint, thereby reducing the necessity for the CNS to exert gating over the somatosensory input. This reduction in proprioceptive feedback might have resulted from the lack of tactile stimulation induced by the splint, a condition known to downregulate proprioceptive feedback^51,52^. Additionally, Golgi tendon organs, which respond to tendon stretch, provide crucial proprioceptive feedback, particularly at the slow speeds required in the present task^49^. The activity of the Golgi organs may have been reduced in the NoTactile group due to decreased friction between the finger and the surface caused by wearing the finger splint. This evidence raises intriguing questions about the potential impact of using a finger splint on the integration of tactile and proprioceptive feedback, warranting further study.

The Tactile group also exhibited inhibition in the source-localized ipsilateral PPC to the drawing hand during mirror drawing. This finding was unexpected and presents a challenge for interpretation. Indeed, the right PPC is crucial for acquiring new visuomotor skills^30,53^ and processing visuospatial information^54,55^. These functions appear particularly relevant for controlling movements in response to mirror-reversed visual feedback. A plausible interpretation for the diminished activity in the right PPC may be related to earlier research indicating that TMS-induced disruption of the right PPC impairs the perception of tactile input in space while leaving proprioceptive judgement unaffected^56^. Thus, the reduced activity of the right PPC in the Mirror vision condition, compared to the Direct vision condition, may represent a decreased reliance on tactile input, which could also enable a more efficient integration of relevant sensory cues (e.g., visual) for controlling the tracing movements.

In an event-related potential study, Bernier et al.^20^ presented evidence that somatosensory gating during tracing with mirror-reversed vision originates from intracortical mechanisms, rather than from peripheral, spinal, or thalamic sources. The PFC, which integrates task requirements^57,58^, is known to exert an inhibitory effect over the primary somatosensory cortex, potentially disrupting the transmission of incoming signals^59,60^. In the present study, the concurrent increased activity in the presumed left PFC, observed alongside somatosensory inhibition in participants of the Tactile group may highlight this role of the PFC in mediating tactile gating. On the other hand, the PPC exhibits dense connections with both the PFC and somatosensory cortex^61^. Therefore, potential explanations for the reduced PPC activity observed in the Tactile group in the Mirror condition include the inhibitory influence of the fronto-parietal network or a decrease in the inflow of tactile sensory information from the somatosensory cortex.

It should be noted that participants in the Tactile group still experienced somatosensory conflict despite evidence of somatosensory gating. This is illustrated by consistently higher jerk index values during the mirror vision condition across the entire experimental session (Supplementary Fig. S1), indicating that the conflict persisted and affected movement smoothness. Consequently, the sensory gating likely reduced, rather than completely eliminated, the processing of somatosensory input. Nevertheless, it can be inferred that without this partial sensory suppression, tracing performance would have deteriorated even further by the mirror-reversed vision.

One could predict increased activation of the visual cortex in both the Tactile and NoTactile groups during mirror tracing. Surprisingly, this effect was exclusively found in the former group, i.e. in participants wearing a finger splint while tracing the shape. Enhancing visual activation and decreasing the activity of the somatosensory cortex are distinct compensatory mechanisms that can both increase the relative weight of visual feedback in the visuo-somatosensory integrative processes. With tactile feedback already reduced due to the finger splint, the NoTactile group may have employed a strategy to reduce sensory conflict by upregulating visual feedback when tracing the shape. This increased reliance on vision aligns with findings from studies involving deafferented patients, which have demonstrated that visual feedback alone is sufficient for controlling movements under mirror-reversed visual conditions^29^. It is also consistent with research showing visual dominance over somatosensory signals in incongruent visuo-somatosensory environments^31,44^. The increased activity was specifically observed in the left visual cortex, which is known to respond more strongly to visual images of the hand compared to the right visual cortex^62,63^. In this vein, the heightened left-lateralized activity observed in the visual cortex may suggest that the hand was preferably represented using visual coordinates during the visuo-somatosensory conflict. The reason why the Tactile group only showed decreased activity of the somatosensory cortex without a significant change in the activity of the visual cortex remains unclear. This observation may suggest difficulty in adopting a sensory weighting strategy that simultaneously modulates visual and somatosensory inputs.

In summary, our findings provide compelling evidence that tactile feedback contributes to controlling the direction of voluntary hand movements when visual cues are present. In an incongruent visuo-somatosensory environment, this tactile feedback increased the experienced sensory conflict, prompting somatosensory gating to reduce the discrepancy. This regulation of tactile feedback suggests a dynamic adjustment in sensorimotor processing to optimize movement execution in challenging perceptual contexts, underscoring the adaptive flexibility of the sensorimotor system.

## Methods

### Experimental protocol

Thirty-two healthy adults (16 men, 16 women, 25±4 years old), with normal or corrected-to-normal vision, participated in the experiment. All participants were right-handed according to the Edinburgh Handedness Inventory (mean laterality score: 82.29 ± 23.6(SD)). All participants gave a fully informed written consent to participate in the study. The number of participants (sample size) was determined by referring to previous studies using the mirror-reversed tracing paradigm^31,48^. The protocols and procedures adhered to the guidelines established in the 1964 Declaration of Helsinki and were approved by the CERSTAPS ethics committee.

Before the experiment, participants washed and dried their hands. Since excessive hydration and lipid concentration of the index finger pulp can alter the friction forces at the skin-surface interface^64^, we characterized the hydrolipid film of each participant using a skin moisture oil content analyzer (Hurrise, China). The measurement was conducted thrice, both prior to and following the experiment, and the average value of each set of 3 recordings was computed. Both groups exhibited normal humidity and oil content values before and after the experiment^65^ (Supplementary Tables S11 & S12). During the experimental session, participants sat comfortably in a dark chamber with their right forearm and elbow resting on a table. Positioned in front of them was a force platform (150 mm x 150 mm, AMTI HE6X6, A-Tech Instruments Ltd., Toronto, Canada), upon which a 3D-printed surface (Sigma D25 printer, BCN3D) made of biopolymer thermoplastic (polylactic acid, PLA; 150 m x 150 m x 2 mm, see Supplementary Fig. S1) was screwed. The surface featured spiral patterns inspired by the sizes and shapes of the dermatoglyphs and of the receptive fields of the type 1 mechanoreceptors (FAI and SAI). This bioinspired surface has been shown to enhance the transmission of cutaneous inputs compared to smooth or grooved surfaces^66,67^. Due to the circular patterns, the spatial frequency of the surface remains virtually constant regardless of the direction of the relative motion between the finger and the surface.

Participants were assigned to Tactile (n=16; 8 women; mean age: 24.3± 3.5 (SD)) and NoTactile (n=16; 8 women; mean age: 24.8± 3.9 (SD)) groups. Data from one participant in the NoTactile group had to be discarded due to technical issues. The task involved tracing the contour of an irregular polygon (100 mm x 80 mm) on the surface. The polygon was composed of 13 thin segments (1 mm wide) of varying lengths, with a total perimeter of 489 mm. Participants in the Tactile group traced the shape using their bare right index finger, while those in the NoTactile group performed the same task wearing a plastic finger splint (BBTO) on their right index finger (see Fig. 1). The measured roughness of the finger splint (i.e., Ra 1.17 µm) was much smaller than the measured roughness of human index fingers (Ra ∼10 µm) (Supplementary Table S13). The finger splint’s smooth surface with its low frictional properties and high hardness (resulting in minimal deformation and thus reduced contact area) created favorable conditions for reducing finger vibration and shear forces, thereby minimizing tactile stimulation during the tracing.

The temporal organization of the trials is presented in Fig. 7. At the beginning of each trial, participants placed their index finger at the shape’s starting point (yellow mark in Fig. 6), adjusting their finger’s normal force in the range between 0.2 N and 0.4 N for 3-5 s (i.e., force normally produced during tactile exploration^68^). The experimenter could see the normal force exerted on the surface in real time and provided feedback to help the participants to comply with this requirement. Then, the participants heard two “beep” signals, randomly interspersed by intervals of 3, 4 or 5 s. The first “beep” indicated the initiation of the static phase during which the participants had to remain motionless without modifying the normal force. The second “beep” marked the onset of the tracing phase, which lasted 11 s and ended by a final tone signal. Participants had no difficulty keeping the normal force relatively constant during the tracing (Supplementary Fig. S5). Maintaining a stable force range was crucial across both experimental groups and visual conditions, as the magnitude of sensory gating can be influenced by the applied forces^69,70^. The experimental session involved a Direct condition (35 trials), where participants had a direct view of their tracing hand, followed by a Mirror condition (35 trials). In the Mirror condition, participants looked the shape and their hand through a round mirror (Comair Cabinet Executive mirror, diameter 280 mm), placed at eye-level to their front left side with a 45-degree angle (see Fig. 1). A black shield obstructed the direct view of the hand and the shape.

**Fig. 6.**
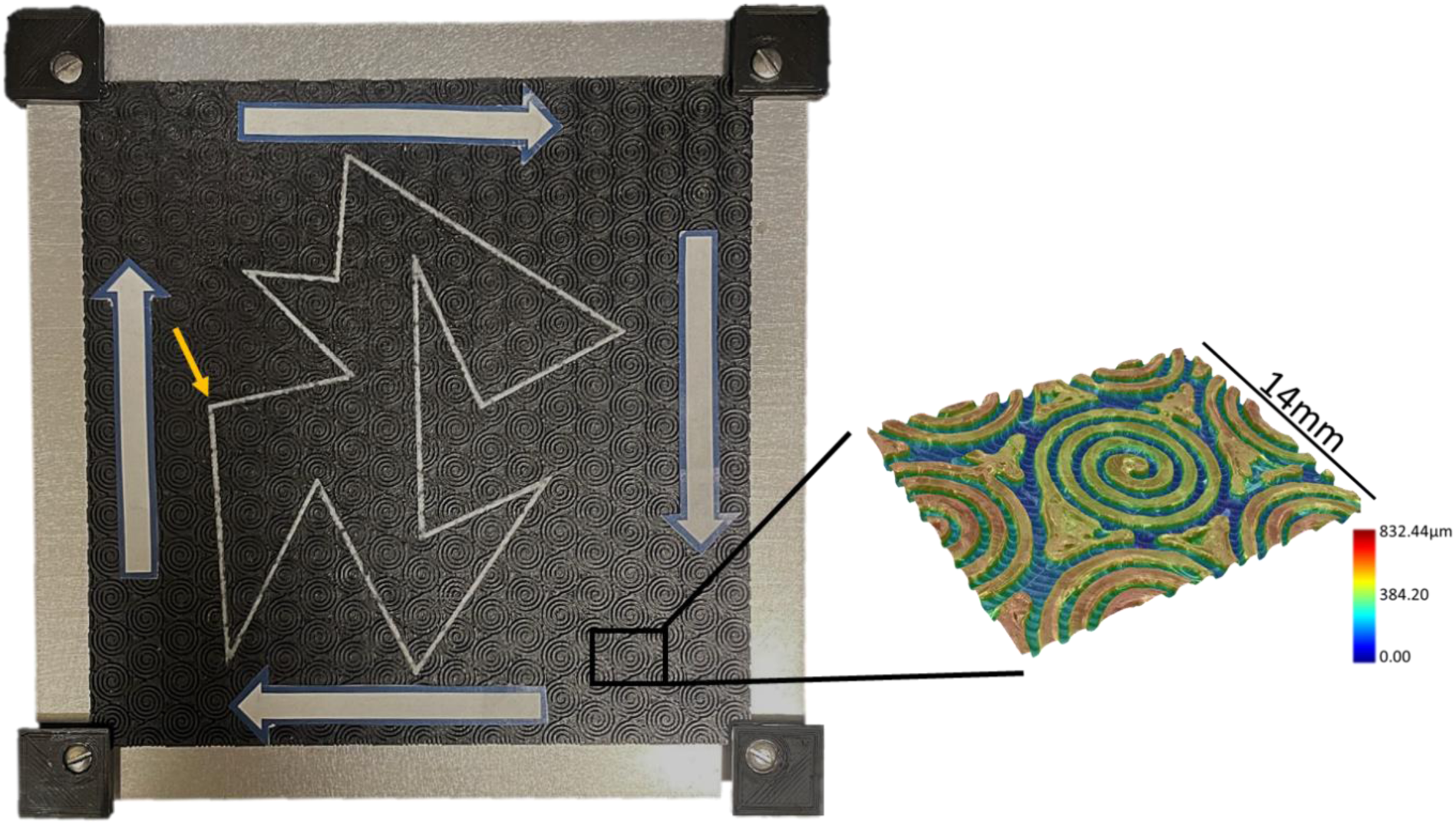
The force platform with the textured surface and the irregular polygon. The finger starting position of the first trial is marked by a yellow arrow (not present during the experiment) and the movement direction is indicated by white arrows affixed on the surface. This information was particularly useful for the participants in the Mirror condition, due to the inverted vision. The starting position for subsequent trials was the ending position of the preceding trials. Right: 3-D image of a sample of the bio-inspired textured surface captured with a VHX-7000 digital microscope. It was made of spirals with a radius of 4 mm, had a spatial periodicity of 0.9 mm, and the depth of the valley between the ridges was 0.7 mm.

**Fig. 7.**
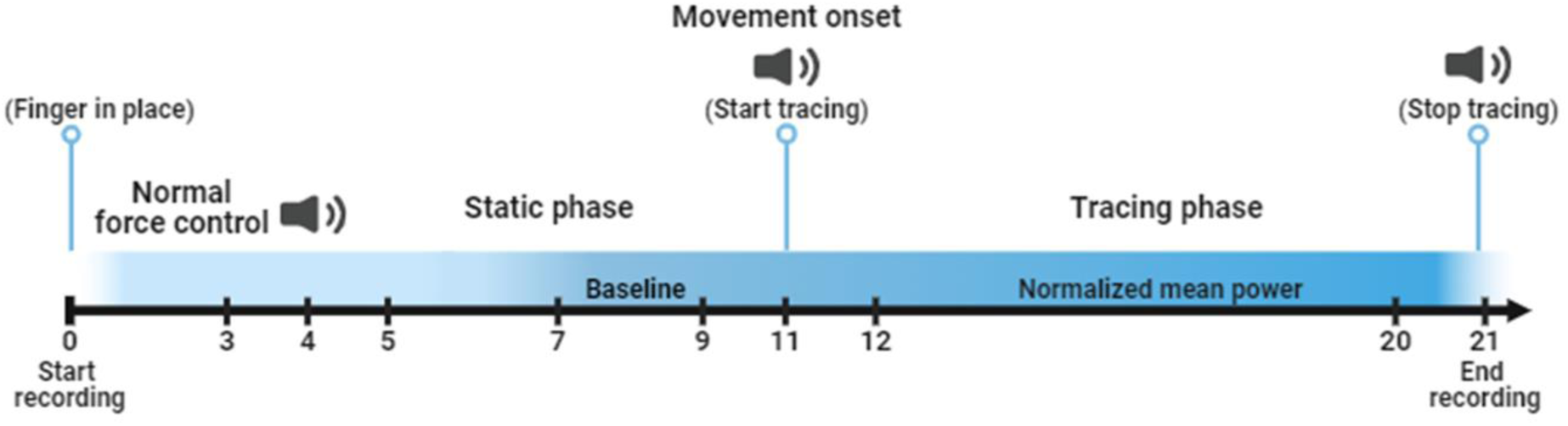
Temporal organization of the trials. The absolute current during the tracing phase was expressed as a change of current amplitude with respect to the mean absolute current computed in a 2-s baseline window ending 2 s before the go signal. Then, we computed the mean absolute current amplitude from 1 to 9 s after the go signal, as indicated by the ‘start tracing’ mark.

When exposed to a sensory conflict, participants’ performance can rapidly return to pre-exposure levels, as evidenced in Sarlegna et al.^71^ where participants regained baseline performance in just 15 trials. Accordingly, our experimental protocol was built with the aim of minimizing participants’ adaptation to their novel sensory environment. This was particularly crucial for ensuring the representativeness of the measured EEG variables, which were obtained by calculating the means of all trials for both visual conditions. First, we designed a shape with several sharp corners which increased the complexity of the tracing in Mirror conditions^45^. The starting position varied across trials, and participants were instructed to move their hand freely while observing it with direct vision every 5 trials. For reasons of homogeneity between the conditions, this procedure was also followed in the Direct condition. Instructions emphasized using slow tracing movements (i.e., ∼10 mm/s as demonstrated by an experimenter before the experiment) to reduce contamination of the EEG signals by fast pursuit eye movements and extensive activation of arm muscles during tracing. In case of deviating from the shape’s outline, participants had to return to the same point where they left the polygon before resuming the tracing process.

## Data Acquisition and Processing

### Behavior

Reaction forces and moments applied by the index finger on the surface were recorded at 256 Hz with the AMTI force platform on which the 2 mm surface was screwed. We used the AMTI’s HE6X6 model, a compact 6-axis force plate capable of measuring loads up to 2.2 N (Fx and Fy) and 4.5 N (Fz) with high-resolution (12-bit). Thus, this model is well suited for capturing the forces applied by the finger while sliding on the surface.

All behavioral variables were computed during the tracing phase, from the occurrence of the “start tracing” signal until the end of the recording (“stop tracing” signal). For both the Tactile and NoTactile groups, tactile stimulation during the tracing was estimated by computing the coefficient of friction, the power spectral density (PSD) of the finger’s vertical acceleration and the amplitude of this acceleration.

The mean friction coefficient was calculated using the following formula on the low pass filtered force data (Butterworth 4th-order, 7 Hz cut-off frequency):

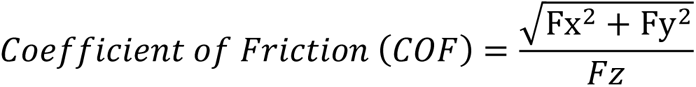

In the NoTactile group, the COF was measured at the interface between the surface and the splint rather between the splint and the index finger. The tangential deformation of the splint (i.e., mechanical stimuli) was then transferred from the splint to the skin. As the splint encompassed a large part of the finger, some of this deformation was transmitted to regions other than the finger pulp. Due to the splint’s smoothness and its stiffness, the tangential deformation of the splint was expected to be minimal.

The vertical acceleration of the right index finger was recorded at 1024 Hz with a small and light (5 mm x 12 mm x 7 mm, 0.8 g) accelerometer (PCB 352A24, PCB Piezotronics, Inc., Depew, USA) secured on the finger’s nail with wax (Fig. 1). With a high sensitivity of 100 mV/g and a measurement range of ±50 g pk, this accelerometer can detect very low acceleration amplitude. Its broadband resolution of 0.0002 g rms further enhances precision, allowing us to capture subtle variations in friction-induced vibrations with remarkable clarity. The broad frequency range of 1 to 8000 Hz ensures coverage of vibration frequencies relevant for the tactile receptors while its lightweight design (0.8 gm) minimizes interference with natural finger movement. To attenuate power line artifacts, a 50 Hz ZapLine filter^72^ was applied to the accelerometer signal. Despite the application of the filter, a small peak at 50 Hz persists, likely attributable to residual electrical noise (Fig. 2a). We computed the power spectral density (PSD) of the vertical acceleration using the Welch method^73^. An average PSD was calculated for every 1-s interval of the recorded signal, with a 50% overlap using Hanning windows. 1024 samples were used for the calculation of the Non-Uniform Fast Fourier Transform (NFFT). We integrated the PSD signal within the 5-45 Hz frequency range, where maximum sensitivity of the direction-sensitive SAI and FAI mechanoreceptors occurs. Additionally, we calculated the amplitude of acceleration by computing the square root of the integral of PSD signal beyond 10 Hz.

The jerk index, estimated here by the derivative of the raw shear forces along the X and Y axes, provides valuable insights into the smoothness and abruptness of force transitions during tracing tasks^36^. It assesses movement quality by revealing the stability and consistency of force application during tracing while remaining sensitive to intermittent disruptions. Tracing performance was assessed by estimating the average sum of absolute jerk index during the defined tracing time window of each trial with the formula:

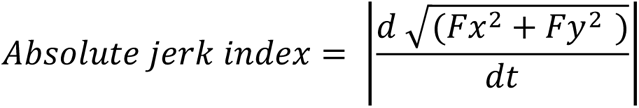

All behavioral data were recorded with a Keithley 12-bit A/D converter device (AD-win pro, Keithely Instruments, Cleveland, OH). Computations were carried out using Matlab 7.0 (The MathWorks, Natick, MA) and custom-made ANALYSE software.

### Electroencephalography (EEG)

EEG signals were continuously recorded using 64 Ag/AgCl electrodes embedded on an elastic cap (BioSemi ActiveTwo system, Amsterdam, The Netherlands). Specific to the Biosemi system, the conventional ground electrode was substituted with a Common Mode Sense (CMS) active electrode and a Driven Right Leg (DRL) passive electrode. The signals were pre-amplified at the electrode sites and further amplified with DC amplifiers. To capture electrooculographic (EOG) activity, 4 Ag/AgCl pre-amplified electrodes were placed near each outer canthus, and above and below the left orbit. The EEG and EOG signals were digitized with a 24-bit resolution and recorded at 1024 Hz using the ActiView acquisition program. EEG pre-processing was performed with BrainVision Analyzer2 software (Brain Products, Gilching, Germany). The signals were referenced to the average activity recorded by all electrodes and segmented for each experimental condition, aligning with tracing initiation. Visual identification of tracing onset relied on detecting the first peak in the coefficient of friction signal following the imperative go signal (i.e., 2^nd^ “beep”). Ocular-related artifact rejection employed the independent component analysis (ICA), while a 50 Hz Notch filter and a DC detrend were applied to mitigate electrical noise. Visual inspection led to the exclusion of epochs with remaining artifacts. Subsequently, an average of 32 ±3(SD) (Direct condition: 32.42 ±3.3(SD) and Mirror condition: 31.9±2.8 (SD)) epochs per participant underwent further analyses.

Cortical sources reconstruction was performed using the Brainstorm software^37^. To resolve the inverse problem, we employed the minimum-norm technique with unconstrained dipole orientations. Forward models were computed with the boundary element method (BEM, 15,002 vertices)^74^ and mapped onto the high-resolution anatomical MRI Colin 27 brain template from the Montreal Neurological Institute (MNI). Opting for enhanced solution precision, we selected a three-layer model for current diffusion comprising the scalp, inner skull, and outer skull, instead of a basic three concentric spheres model^75^.

The current amplitude computed during the tracing phase was normalized with respect to the mean current amplitude during the static baseline period (-4 to -2 s from movement onset). These normalized values were then averaged across all trials within each group and condition, specifically during the 1-9 s period following the movement onset.

### Statistical Analysis

We used STATISTICA 8.0 (StatSoft, Inc., USA) to analyze the data, with complete details of all statistical analyses provided in the supplementary materials (Tables S1 to S9). The normal distribution of data sets was assessed using the Shapiro–Wilk test to validate the use of parametric tests. When necessary, logarithmic transformation was applied to meet the assumptions of the ANOVA model. All behavioral variables were submitted to 2 (Group: Tactile, NoTactile) × 2 (Vision: Direct, Mirror) mixed ANOVAs (alpha level set at 0.05), with Vision as a repeated factor. Effect sizes are reported as partial eta-squared (η²) values for the ANOVAs. Significant effects were further analyzed using Newman-Keuls post-hoc tests (alpha level set at 0.05). The results are presented using violin plots, which offer a comprehensive views of data distributions, including individual data points, as well as the median and mean.

Within each group, we estimated the effect of the visuo-somatosensory conflict on the topography and amplitude of neuronal activity by contrasting the estimated sources of the absolute EEG current in the Direct and Mirror conditions using t-tests, with a significance threshold set at p < 0.05 (uncorrected). We specifically compared the impact of mirror-reversed vision on the activity of the presumed somatosensory cortex between the Tactile and Notactile groups. This comparison was done by computing the mean absolute current amplitude for each participant within a region of interest (ROI) encompassing the left postcentral gyrus as defined by Destrieux et al.^39^. These mean values were then subjected to a 2 (Group: Tactile, NoTactile) x 2 (Vision: Direct, Mirror) ANOVA with Vision as the repeated factor. The results of these analyses are presented using violin plots.

## Acknowledgements

The project leading to this publication has received funding from the French government under the “France 2030” investment plan managed by the French National Research Agency (reference: ANR-16-CONV000X / ANR-17-EURE-0029), from Excellence Initiative of AixMarseille University - A*MIDEX (AMX-19-IET-004) and from the ANR COMTACT (ANR 2020-CE28-0010-03). We thank Franck Buloup for developing the software Docometre used for data acquisition, Marcel Kaszap for developing the software Analyse used for data processing and all the participants for taking part in the study.

## Author contributions

J.B., M-E.V., and L.M. conceived and designed the study. D.P. contributed to surface fabrication and accelerometer measurements. M-E.V., J.L., and C.S. collected the data. M-E.V., M.S., F.M. analyzed the data. M-E.V. drafted the manuscript. M-E.V., J.B, L.M., M.S., and F.M. reviewed and edited the manuscript.

