## Supplementary for "Guided by touch: Tactile Cues in Hand Movement Control"

### Supplementary material

| Effect | SS | Degrees of Freedom | MS | F | p |
|---|---|---|---|---|---|
| **Intercept** | 1098.62 | 1 | 1098.62 | 3924.13 | 0.00 |
| **Group** | 3.18 | 1 | 3.18 | 11.35 | 0.002 |
| **Error** | 7.56 | 27 | 0.28 | | |
| **Visual condition** | 0.03 | 1 | 0.03 | 0.12 | 0.73 |
| **Visual condition * Group** | 0.03 | 1 | 0.03 | 0.15 | 0.71 |
| **Error** | 5.37 | 27 | 0.2 | | |

**Table S1.** Results of the 2X2 ANOVA analysis on the logarithmic data of the integrated PSD accelerometer signal for the 5-45 Hz range. A significant Group effect was yielded, indicating statistically significant differences between the Tactile and the NoTactile groups in the accelerometer power within this frequency range. However, there was no significant main effect of Visual condition, nor was there a significant interaction effect between Visual condition and Group. These results suggest that the differences observed in the integrated PSD accelerometer signal are primarily influenced by the experimental grouping rather than the visual condition.

| Effect | SS | Degrees of Freedom | MS | F | p |
|---|---|---|---|---|---|
| **Intercept** | 221.69 | 1 | 221.69 | 3484.6 | 0.00 |
| **Group** | 0.41 | 1 | 0.41 | 6.38 | 0.017 |
| **Error** | 1.71 | 27 | 0.06 | | |
| **Visual condition** | 0.00 | 1 | 0.00 | 0.00 | 0.99 |
| **Visual condition * Group** | 0.01 | 1 | 0.01 | 0.27 | 0.6 |
| **Error** | 0.99 | 27 | 0.04 | | |

**Table S2.** The results of the 2X2 ANOVA analysis on the logarithmic data of the square root value of the integrated PSD accelerometer signal above 10 Hz revealed a significant effect for the Group factor, indicating differences between the two groups.

| Effect | SS | Degrees of Freedom | MS | F | p |
|---|---|---|---|---|---|
| **Intercept** | 7.18 | 1 | 7.18 | 195.52 | 0.0000 |
| **Group** | 3.08 | 1 | 3.08 | 83.93 | 0.0000 |
| **Error** | 1.06 | 29 | 0.037 | | |
| **Visual condition** | 0.001 | 1 | 0.001 | 0.51 | 0.48 |
| **Visual condition * Group** | 0.0001 | 1 | 0.0001 | 0.07 | 0.8 |
| **Error** | 0.06 | 29 | 0.002 | | |

**Table S3.** The results of the 2X2 ANOVA analysis on the logarithmic data of the coefficient of friction yielded a significant effect of the Group factor, demonstrating that friction levels significantly differed between the groups. However, there was no significant effect of the Visual condition factor, suggesting that the visual condition did not influence the friction levels.

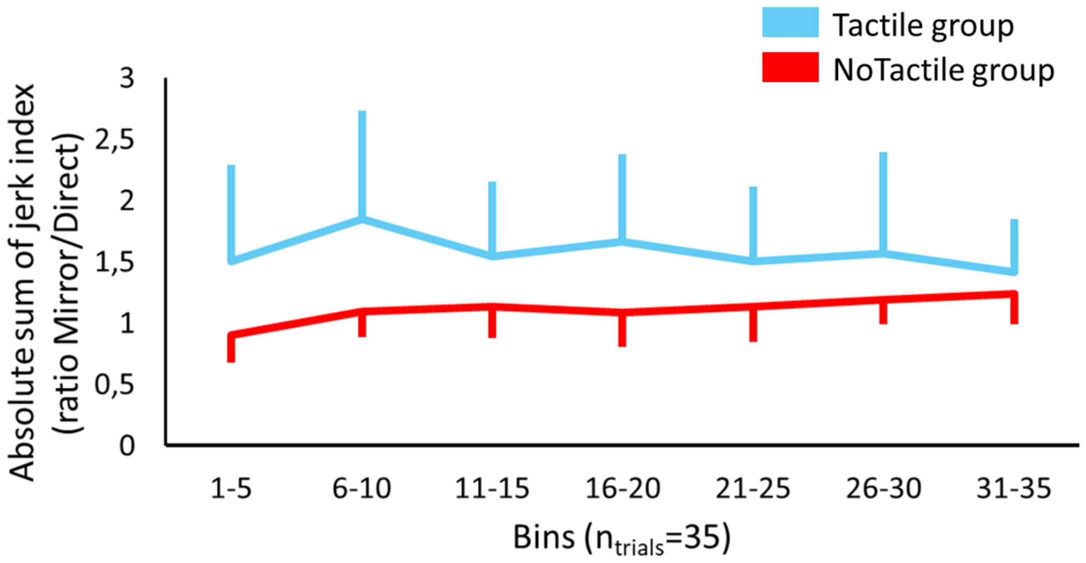


**Fig. S1 |** The evolution of the Mirror/Direct ratio of the sum of absolute jerk index over the course of 35 trials for the Tactile (blue) and the NoTactile group (red), pooled in 7 bins of 5 consecutive trials to control for adaptation to the sensory conflict. The jerk values computed in the Mirror condition were normalized with respect to the jerk values computed in the Direct condition. A 2 (Group: Tactile. NoTactile) × 7 (Bin: $Bin_{1\text{-}5}$, $Bin_{6\text{-}10}$, $Bin_{11\text{-}15}$, $Bin_{16\text{-}20}$, $Bin_{21\text{-}25}$, $Bin_{26\text{-}30}$, $Bin_{31\text{-}35}$) mixed ANOVA analysis revealed a significant effect of the experimental group ($F_{1,25}$=16.47, p=0.01, partial $\eta^2$=0.24). Neither the Bin factor nor the Group*Bin interaction yielded a significant effect. The fact that the jerk index remained relatively constant across the 35 trials for each group provides behavioral bases for comparing, between the groups, the averaged EEG activity computed from all trials.

| Effect | SS | Degrees of Freedom | MS | F | p |
|---|---|---|---|---|---|
| **Intercept** | 1.33 | 1 | 1.33 | 16.47 | 0.0004 |
| **Group** | 0.64 | 1 | 0.64 | 7.87 | 0.01 |
| **Error** | 2.02 | 25 | 0.08 | | |
| **Bin** | 0.12 | 6 | 0.02 | 1.94 | 0.08 |
| **Bin * Group** | 0.08 | 6 | 1.38 | 1.39 | 0.22 |
| **Error** | 1.49 | 150 | | | |

**Table S4.** Results of the 2X7 ANOVA analysis on the logarithmic data of the absolute sum of jerk for the factor Bin. The results showed significant effect only for the Group factor. The effects of Bin and the interaction between Bin and Group were not significant, indicating that the observed differences in jerk index between the groups were not affected by the stage of the task.

| Effect | SS | Degrees of Freedom | MS | F | p |
|---|---|---|---|---|---|
| **Intercept** | 10.91 | 1 | 10.91 | 178.49 | 0.000 |
| **Group** | 1.02 | 1 | 1.02 | 16.75 | 0.0003 |
| **Error** | 1.77 | 29 | 0.06 | | |
| **Vision** | 0.14 | 1 | 0.14 | 8.17 | 0.008 |
| **Vision * Group** | 0.11 | 1 | 0.11 | 6.45 | 0.017 |
| **Error** | 0.5 | 29 | 0.02 | | |

| Cell number | Group | Visual condition | {1}<br>0.46 | {2}<br>0.64 | {3}<br>0.29 | {4}<br>0.3 |
|---|---|---|---|---|---|---|
| **1** | Tactile | Direct | | 0.0008 | 0.05 | 0.03 |
| **2** | Tactile | Mirror | 0.0008 | | 0.0002 | 0.0001 |
| **3** | NoTactile | Direct | 0.05 | 0.0002 | | 0.82 |
| **4** | NoTactile | Mirror | 0.03 | 0.0001 | 0.82 | |

**Table S5 & S6.** Results of the 2X2 ANOVA analysis and the Newman-Keuls post-hoc test on the average absolute sum of jerk index data. Pooled MSE=0.039, df=44.01. Significant main effects for both Group and Vision were yielded, indicating that there are differences in the average absolute sum of jerk index between the experimental groups and visual conditions. The significant Vision*Group interaction revealed that the jerk index significantly increased during mirror tracing only in the Tactile group, indicating performance deterioration. In contrast, the NoTactile group showed no significant changes in jerk index between mirror and direct conditions.

| Effect | SS | Degrees of Freedom | MS | F | p |
|---|---|---|---|---|---|
| **Intercept** | 1.58 | 1 | 1.58 | 841.69 | 0.00 |
| **Group** | 0.009 | 1 | 0.09 | 4.78 | 0.038 |
| **Error** | 0.05 | 27 | 0.002 | | |
| **Visual condition** | 0.01 | 1 | 0.01 | 3.18 | 0.09 |
| **Visual condition * Group** | 0.002 | 1 | 0.002 | 4.6 | 0.04 |
| **Error** | 0.009 | 27 | 0.0003 | | |

| Cell number | Group | Visual condition | {1}<br>0.16 | {2}<br>0.14 | {3}<br>0.18 | {4}<br>0.18 |
|---|---|---|---|---|---|---|
| **1** | Tactile | Direct | | 0.01 | 0.25 | 0.39 |
| **2** | Tactile | Mirror | 0.01 | | 0.03 | 0.03 |
| **3** | NoTactile | Direct | 0.25 | 0.03 | | 0.8 |
| **4** | NoTactile | Mirror | 0.39 | 0.03 | 0.8 | |

**Table S7 & S8.** Results of the 2X2 ANOVA analysis and the Newman-Keuls post-hoc test on the mean absolute current amplitude over the left postcentral gyrus ROI. Pooled MSE=0.001, df=36.32. A statistically significant effect was revealed for the factor Group but not for the factor Visual condition, showing different activation levels over the defined ROI between the two groups. A significant interaction effect between Visual condition and Group was also yielded and post-hoc comparisons indicate participants in the Tactile group exhibited significantly different somatosensory responses between the Direct and Mirror conditions, while those in the NoTactile group did not show such a difference.

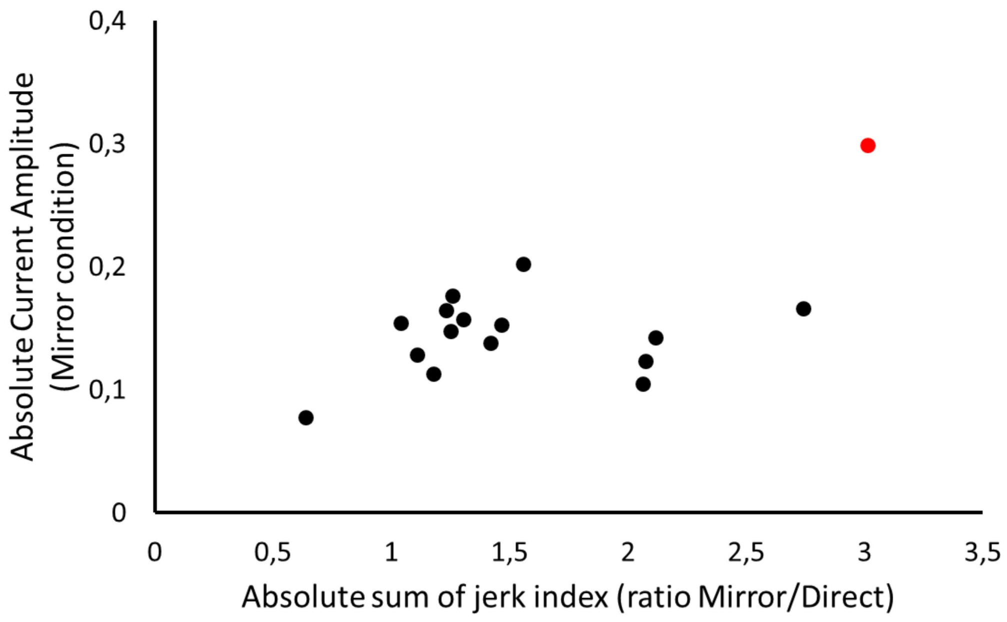


**Fig. S2 |** Scatter plot illustrating the Mirror to Direct ratio of the absolute sum of jerk index over the absolute current amplitude during the Mirror condition for the Tactile group. Each data point represents an individual participant, with the red dot indicating an outlier. This participant displayed both the highest absolute current amplitude and the greatest Mirror/Direct ratio of jerk index value among all participants in the Tactile group.

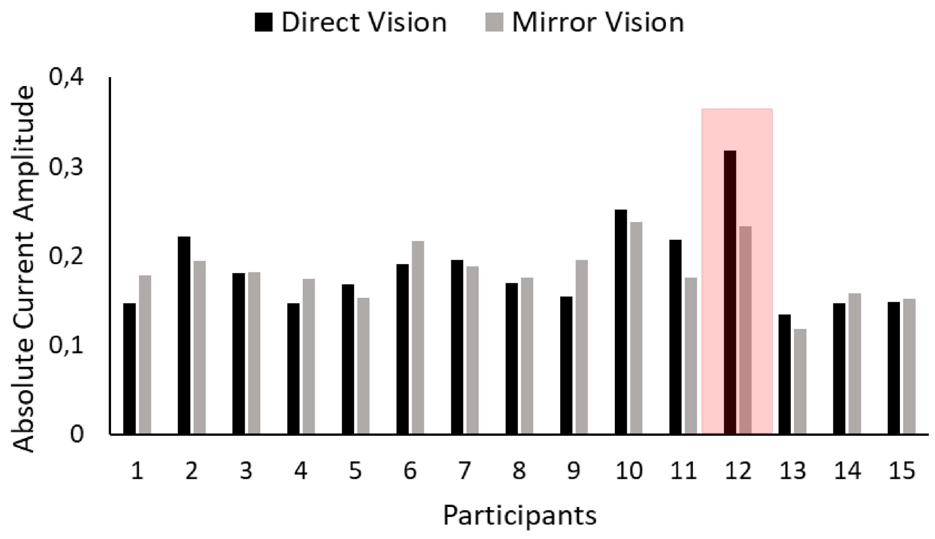


**Fig. S3 |** Bar graph of the individual absolute current amplitude during the Direct Vision (black) and Mirror Vision (grey) condition. The red frame highlights the outlier data, depicting the greatest decrease in current amplitude from the Direct to the Mirror condition among all participants within the NoTactile group.

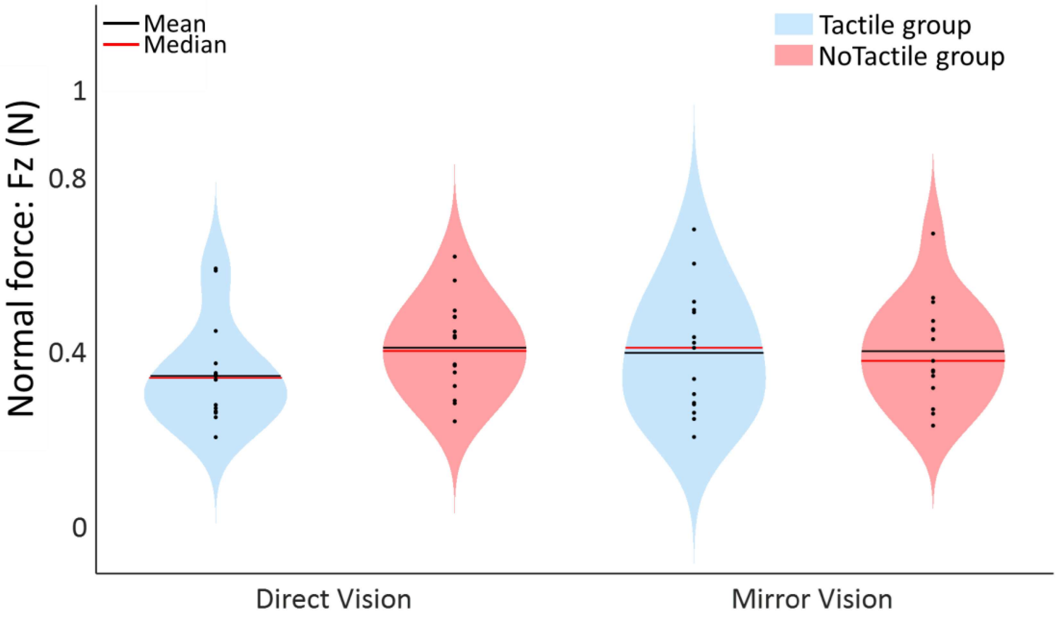


**Fig. S4 | Normal force remains consistent across both experimental groups and conditions.** Violin plots of the mean normal force of all the trials for the Tactile group (blue) and the NoTactile group (red) during Direct and Mirror tracing. Individual data points are represented with scatter plots within each violin plot.

| Effect | SS | Degrees of Freedom | MS | F | p |
|---|---|---|---|---|---|
| **Intercept** | 9.69 | 1 | 9.69 | 416.86 | 0.00 |
| **Group** | 0.008 | 1 | 0.008 | 0.34 | 0.57 |
| **Error** | 0.67 | 29 | 0.02 | | |
| **Visual condition** | 0.02 | 1 | 0.02 | 3.62 | 0.067 |
| **Visual condition * Group** | 0.01 | 1 | 0.01 | 2.85 | 0.10 |
| **Error** | 0.14 | 29 | 0.005 | | |

**Table S9**. Results of the 2X2 ANOVA analysis on the logarithmic data of the normal force. No significant effects for either Group or the Visual condition factor, nor for the interaction effect between Visual condition and Group were revealed, indicating that irrespective of the visual condition, both experimental groups maintained a consistent range of normal force applied on the platform throughout the experiment.

| Tactile group | Humidity content (a.u) Before | Humidity content (a.u) After | Oil content (a.u.) Before | Oil content (a.u.) After |
|---|---|---|---|---|
| S1_F | 56.2 | 60.1 | 23.6 | 27 |
| S2_F | 50.3 | 64.57 | 22.47 | 29 |
| S3_F | 52.4 | 64.1 | 23.5 | 28.8 |
| S4_F | 50.3 | 59 | 22.7 | 26.5 |
| S5_F | 56.23 | 55.1 | 25.27 | 24.77 |
| S6_F | 53.1 | 57.97 | 23.9 | 24.03 |
| S7_F | 57.17 | 59.17 | 22.97 | 26.6 |
| S8_F | 55.37 | 58.2 | 24.87 | 27.6 |
| S1_M | 56.3 | 55.47 | 25.3 | 26.4 |
| S2_M | 59.57 | 61.93 | 26.73 | 27.8 |
| S3_M | 61.57 | 64.3 | 27.67 | 29.8 |
| S4_M | 50.33 | 53.73 | 22.6 | 24.13 |
| S5_M | 61.17 | 58.87 | 27.47 | 27.1 |
| S6_M | 60.3 | 73.93 | 27.1 | 33.1 |
| S7_M | 44.13 | 54.93 | 19.8 | 24.7 |
| S8_M | 57.1 | 61.83 | 25.67 | 27.8 |

| NoTactile group | Humidity content (a.u) Before | Humidity content (a.u) After | Oil content (a.u.) Before | Oil content (a.u.) After |
|---|---|---|---|---|
| S1_F | 61.27 | 63.07 | 27.53 | 28.33 |
| S2_F | 58.6 | 61.13 | 26.3 | 27.47 |
| S3_F | 46.73 | 53.2 | 20.9 | 23.9 |
| S4_F | 60.7 | 62.6 | 27.27 | 28.1 |
| S5_F | 59.6 | 63.4 | 26.8 | 28.5 |
| S6_F | 60.17 | 61.3 | 27 | 27.53 |
| S7_F | 53.77 | 57.22 | 24.13 | 25.1 |
| S8_F | 57.67 | 71.03 | 25.77 | 31.93 |
| S1_M | 59.37 | 64.6 | 26.7 | 29.03 |
| S2_M | 61.33 | 65.9 | 27.57 | 28.7 |
| S3_M | 62.57 | 55.13 | 28.1 | 24.73 |
| S4_M | 48.3 | 51.37 | 21.67 | 23.07 |
| S5_M | 55.23 | 57.3 | 24.8 | 26.2 |
| S6_M | 52.57 | 52.37 | 23.63 | 23.5 |
| S7_M | 50.1 | 59.33 | 22.5 | 26.63 |

**Table S11 & S12.** Index finger composition, measured before and after the experimental session using a skin analyzer, and characterized by the humidity and oil content in arbitrary units for the Tactile (Left) and the NoTactile (Right) groups.

| Roughness sample | Arithmetic Average Roughness – Ra (µm) | Mean Roughness Depth – Rz (µm) |
|---|---|---|
| Finger splint | 1.17 | 6.5 |
| Male fingertip | 10.45 | 41.96 |
| Female fingertip | 9.5 | 35.37 |

**Table S13.** Values of the sample's roughness as measured using a stylus profilometer (MarSurf PS 10) with Inductive skidded probe (resolution 8 nm). The roughness of the fingertips was obtained using a resin mold of the operator's fingertips. Ra is a measure of the average deviation of the surface valleys and peaks from the mean line of the surface within a sampling length, thus serves as a good indicator of the overall roughness of a sample. Rz indicates the vertical distance between the highest peak and the lowest valley within the sampling length, giving insight into the height of the surface irregularities. The values are obtained by averaging the Ra and Rt values obtained on 6 different profiles measured along perpendicular directions (3 lines par direction) on the fingertip contact zone.
